# Integrated Dual-Channel Retrograde Signaling Directs Stress Responses by Degrading the HAT1/TPL/IMPα-9 Suppressor Complex and Activating CAMTA3

**DOI:** 10.1101/2024.08.29.610327

**Authors:** Liping Zeng, Jingzhe Guo, Malathy Palayam, Carlos Rodriguez, Maria Fernanda Gomez Mendez, Yaqi Wang, Wilhelmina van de Ven, Jose Pruneda-Paz, Nitzan Shabek, Katayoon Dehesh

**Author notes:** Corresponding author: Katayoon Dehesh, Address: Genomics Building 4119C, University of California, Riverside, Riverside, CA 92521.

## Abstract

The intricate communication between plastids and the nucleus, shaping stress-responsive gene expression, has long intrigued researchers. This study combines genetics, biochemical analysis, cellular biology, and protein modeling to uncover how the plastidial metabolite MEcPP activates the stress-response regulatory hub known as the Rapid Stress Response Element (RSRE). Specifically, we identify the HAT1/TPL/IMPα- 9 suppressor complex, where HAT1 directly binds to RSRE and its activator, CAMTA3, masking RSRE and sequestering the activator. Stress-induced MEcPP disrupts this complex, exposing RSRE and releasing CAMTA3, while enhancing Ca^2+^ influx and raising nuclear Ca^2+^levels crucial for CAMTA3 activation and the initiation of RSRE- containing gene transcription. This coordinated breakdown of the suppressor complex and activation of the activator highlights the dual-channel role of MEcPP in plastid-to-nucleus signaling. It further signifies how this metabolite transcends its expected biochemical role, emerging as a crucial initiator of harmonious signaling cascades essential for maintaining cellular homeostasis under stress.

**Summary:** This study uncovers how the stress-induced signaling metabolite MEcPP disrupts the HAT1/TPL/IMPα-9 suppressor complex, liberating the activator CAMTA3 and enabling Ca^2+^ influx essential for CAMTA3 activation, thus orchestrating stress responses via repressor degradation and activator induction.

## Introduction

In the intricate interplay of dynamic environmental cues and the regulation of growth and stress responses, plants rely on finely orchestrated intraorganellar communications. These communications are facilitated, in part, by metabolites that serve dual roles in both biochemical pathways and signaling networks. Among these metabolites, 2-C-methyl-d-erythritol-2,4-cyclopyrophosphate (MEcPP) emerges as a pivotal intermediate in the plastidial methylerythritol phosphate (MEP) pathway. Beyond its role as a precursor for isoprenoids, MEcPP also assumes significance as a stress-specific retrograde signaling metabolite^1^.

Elevated levels of MEcPP triggered by stress stimuli within plastids initiate a complex signaling cascade, orchestrating a series of events that ultimately lead to the expression of stress-response nuclear genes. Previous studies have shown that elevated MEcPP levels trigger a cascade of events culminating in the activation of calcium-dependent calmodulin-binding transcription activators 3 (CAMTA3)^2^. Subsequently, CAMTA3 activates the Rapid Stress Response Element (RSRE; CGCGTT), a regulatory motif prevalent in about 30% of stress-responsive genes and notably enriched in MEcPP- induced genes^2^. This led to the notion of MEcPP function as a calcium rheostat, ultimately activating stress-responsive genes and linking plastidial stress perception to nuclear gene expression^2^. It’s noteworthy that CAMTA3, despite its reported role as an activator in cold-response, general stress response, and glucosinolate metabolism regulation^2–4^, has also been identified as a transcriptional repressor for immunity-related genes^5^. This delicate equilibrium in transcriptional regulation, toggling between gene repression and activation, mirrors observations in the homeodomain-leucine zipper (HD-Zip) protein family, encompassing HD-Zip I-IV groups pivotal for plant responses to environmental cues^6,7^. Notably, HAT1, also known as JAIBA, a member of the class II HD-Zip proteins, showcases a multifaceted nature by acting as an activator of transcriptional pathways in plant development^8^. Simultaneously, it functions as a repressor, inhibiting anthocyanin, auxin, brassinosteroid, ABA, and gibberellin pathways^9–13^, partly through the recruitment of transcriptional co-repressors such as the TOPLESS (TPL) protein, facilitated by its N-terminal ERF-associated amphiphilic repression (EAR) motif^13^.

In Arabidopsis, the TPL/TPR family is categorized into two distinct groups based on sequence similarities, reflecting their evolutionary divergence and specific roles. These proteins are instrumental in governing developmental pathways, mediating hormone signaling, and orchestrating responses to environmental stresses^14,15^. By engaging with diverse transcription factors at gene promoter sites, the TPL/TPR proteins intricately regulate plant growth and enhance adaptability to environmental cues^16,17^.

Eukaryotic cells rely on the nuclear membrane system to meticulously manage transcription and translation processes, ensuring precise regulation. Central to this system is the nuclear import machinery, which facilitates the transportation of proteins, including transcriptional regulators, into the nucleus. These regulators play pivotal roles in governing various aspects of plant growth, development, and responses to environmental stimuli^18–20^. Within this intricate network, nuclear importins, comprising importin α and β, assume key functions in orchestrating responses to diverse stress conditions^21^. Recently, we identified IMPα-9 as a critical component of the MEcPP-mediated plastidial retrograde signaling pathway^22^. Stress-induced accumulation of MEcPP triggers the proteasome system to degrade IMPα-9 and its associated proteins, thereby alleviating their suppression of stress-response genes. This suppression mechanism is part of a broader regulatory network, but direct evidence elucidating how IMPα-9 and its interactors suppress stress-response genes remains elusive.

In this study, we establish the dual functionality of HAT1. First, it directly binds to the RSRE motif and interacts with its activator, CAMTA3. Second, we confirm HAT1’s role as a suppressor of RSRE by forming a complex with TPL/IMPα-9. This complex effectively blocks RSRE’s access to its activator, CAMTA3. However, MEcPP disrupts this complex, exposing RSRE, releasing CAMTA3, which is activated by the increase in intracellular calcium levels. This cascade ultimately triggers the expression of stress-response genes. Our findings highlight the multifaceted and integrated roles of plastidial retrograde signaling in maintaining a delicate balance between transcriptional activators and repressors, crucial for shaping the dynamic landscape of gene expression.

## Results

### HAT1 binds to and suppresses RSRE induction

Our previous research identified RSRE in 30% of stress-induced genes, many of which are activated by elevated MEcPP levels during stress^2^. To gain deeper insight into the RSRE-regulated transcriptional network, we conducted a yeast one-hybrid (Y1H) assay and identified HAT1 as a primary candidate transcriptional regulator binding to RSRE. This finding was supported by both lacZ and luciferase reporter systems (Fig. 1a-1b).

**Figure 1.**
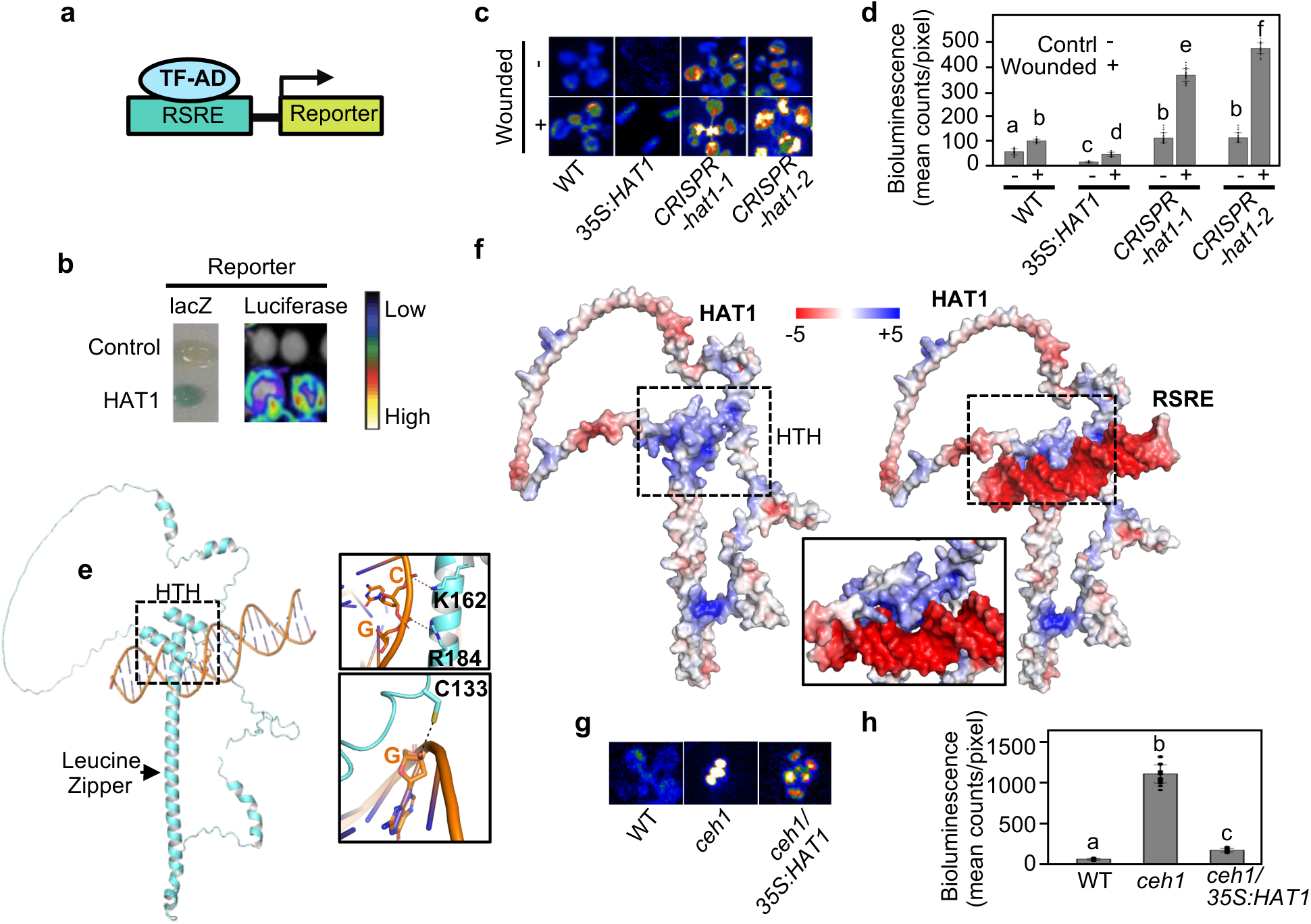
HAT1 binds to and suppresses RSRE induction. **a-b,** RSRE fused to reporter construct used in yeast-one-hybrid using RSRE::LACZ reporter and RSRE:: luciferase construct in *Nicotiana benthamiana* transient assay show HAT1 binding to the RSRE motif. **c-d,** Representative dark-field images of RSRE::Luciferase activity in wild type (WT), HAT1 overexpression (35S:*HAT1*), *CRISPR-hat1-1*, and *CRISPR-hat1-2* lines under unwounded (-) and 90 minutes post- wounding (+) conditions, and **d,** the quantitative measurements of luciferase activities. **e,** AlphaFold 3 structure of HAT1 (cyan)-RSRE (orange) complex (left panel). Zoom-in view of hydrogen bonding between HAT1 amino acids and RSRE (right panel). **f,** Calculated electrostatic surface charge representation of HAT1 (left panel) and HAT1- RSRE complex (right panel). Arrow represents the positively charged region of Helix- turn-Helix (HTH) of HAT1 (left). Dashed box on the right represents the negatively charged RSRE complex (right panel). Zoom in view of the association between HAT1- RSRE complex (middle panel). The structural analysis was based on highest probability model predicted by AlphaFold 3. The surface charge distribution is calculated using the ABPS program implemented in PyMOL. **g-h,** Representative dark-field images of RSRE::Luciferase activity in wild type (WT), *ceh1*, HAT1 overexpression in the ceh1 background (*ceh1/35S*), and their quantitative measurements of luciferase activities in the corresponding lines. Bars that do not share a letter represent statistically significant differences (p<0.05) by ANOVA test with Tukey’s honest significant difference (HSD) test. The error bar is the standard deviation of biological replicates. The color-coded bar displays the intensity of LUC activity.

To further investigate HAT1’s biological role in in-planta transcriptional regulation of RSRE, we manipulated *HAT1* expression levels genetically, generating both overexpression and knockdown lines. In the overexpression lines, we introduced *35S:HAT1* into the background of the 4xRSRE:Luciferase (LUC) construct^23^ (referred to as *35S:HAT1*). For knockdown studies, we utilized the CRISPR-Cas9 system to generate *HAT1* knockout lines, designated as *CRISPR-hat1-1* and *CRISPR-hat1-2* (Extended Data Fig. 1). Our analysis of RSRE-driven LUC activity in the control *4xRSRE::LUC/Col-0* (referred to as WT), *35S:HAT1*, and *CRISPR-hat1* lines revealed diminished bioluminescence signals in the *35S:HAT1* line compared to the two *CRISPR-hat1* lines, and conversely, the *CRISPR-hat1* lines exhibited heightened bioluminescence signals under both standard and wounded conditions compared to the control (Fig. 1c-1d). These findings solidify HAT1’s role as an in-planta suppressor of RSRE activation. Utilizing AlphaFold 3^24^ we accurately predicted the 3D molecular architecture of the HAT1-RSRE complex with high confidence (Fig. 1e-1f). The structure of HAT1 reveals distinct domains, including an N-terminal homeodomain and a HD-Zip region. Notably, the HD-Zip region is characterized by a helix-turn-helix (HTH) motif, enriched with basic residues. These electrostatic properties, conserved across land plants (Extended Data Fig. 2), contribute to the stable formation of the HTH-RSRE complex (Fig. 1e).

These findings prompted us to investigate HAT1’s ability to suppress the previously reported MEcPP-inducible RSRE activity^2^. Thus, we introduced the *35S:HAT1* vector into the constitutively high MEcPP accumulating mutant line, *ceh1*, and generated the *ceh1/35S:HAT1* plants. Subsequent luciferase activity assays revealed a substantial reduction in the constitutive luciferase activity in the *ceh1/35S:HAT1* line compared to the *ceh1* background (Fig. 1g-1h). These results collectively demonstrated that HAT1 acts as a suppressor of MEcPP-induced RSRE activation.

In summary, our findings reveal HAT1’s ability to bind to the RSRE *cis*-element and attenuate MEcPP-induced RSRE activation.

### MEcPP regulation of *HAT1* expression is auxin-dependent

Given the MEcPP-induced RSRE response, we explored a potential feedback loop in which elevated MEcPP levels interfere with HAT1 function through various mechanisms, including transcriptional regulation. This proposition is supported by MEcPP’s ability to reshape the transcriptional landscape. To validate this hypothesis, we initially examined *HAT1* expression levels in the *ceh1* mutant compared to WT plants (Fig. 2a). The reduced expression of *HAT1* in *ceh1* compared to WT plants prompted us to investigate its expression levels in response to wounding, a stress known to induce MEcPP accumulation^1^, a phenomenon reaffirmed in this study (Fig. 2b). The results demonstrate a decrease in *HAT1* transcript levels following wounding (Fig. 2c). These collective findings lend support to the concept of MEcPP-mediated transcriptional suppression of *HAT1*.

**Figure 2.**
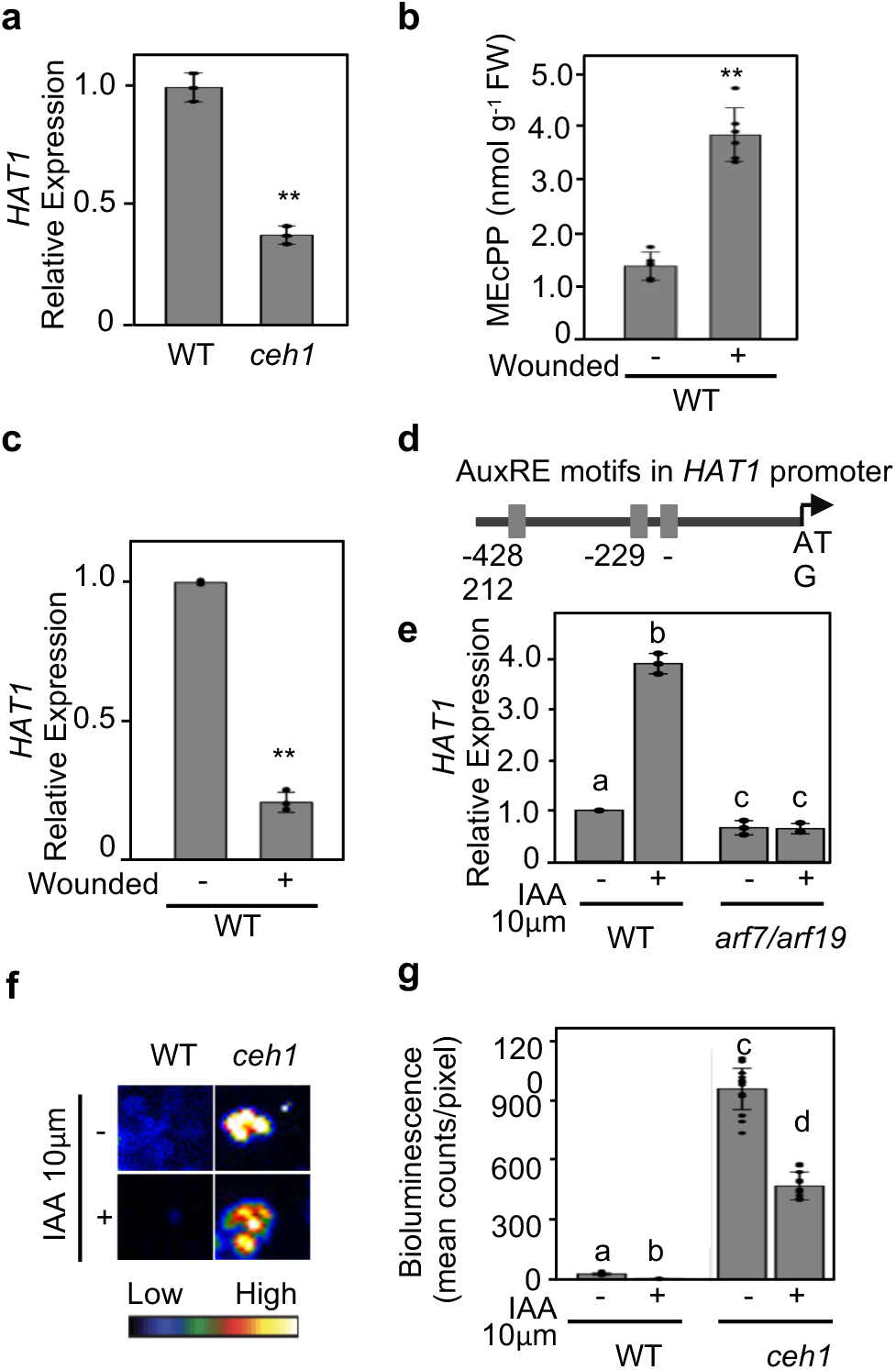
MEcPP regulation of *HAT1* expression is auxin-dependent. **a,** The transcript levels of *HAT1* are reduced in the high MEcPP-containing *ceh1* mutant. **b,** MEcPP levels in leaves of unwounded and 90 minutes post-wounding leaves per fresh weight (FW) unit. **c,** Relative transcript levels of *HAT1* in 90 minutes post-wounded (+) compared to the unwounded control (-). **d,** The schematic presentation of the *HAT1* promoter noting the positions of auxin response elements (AuxRE). **e,** The transcript levels of *HAT1* are induced post 30 minutes 10 μM IAA treatment (+) and are highly diminished in *arf7/arf19* double mutant plants. The transcript level data are presented as the mean value ± SD from three independent biological replicates, each with three technical replicates. Lowercase letters above the bars indicate statistically significant differences (P<0.05) determined by ANOVA test with Tukey’s honest significant difference (HSD) test. The error bar represents the standard deviation of biological replicates. **f-g,** Representative dark-field images of RSRE::Luciferase activity in wild type (WT/*RSRE*) and *ceh1*/*RSRE* plants post 1h 10 μM IAA (+) and mock (-) treatment, and g, quantitative measurements of luciferase activities. The color-coded bar displays the intensity of LUC activity. Lowercase letters above the bars indicate statistically significant differences (P<0.05) determined by ANOVA test with Tukey’s honest significant difference (HSD) test. The error bar represents the standard deviation of biological replicates.

In exploring the regulatory elements governing *HAT1* transcription, we analyzed its promoter sequence and identified three auxin response *cis*-elements (AuxRE) (Fig. 2d). This discovery prompted us to assess *HAT1* transcript levels in both mock-treated and IAA-treated WT plants. We observed a significant increase in *HAT1* transcript levels 30 minutes post IAA treatment, compared to untreated plants, underscoring the auxin-dependent regulation of *HAT1* expression (Fig. 2e). Moreover, considering MEcPP’s established role in diminishing auxin levels^25^ and influencing the transcript levels of ARF7 and ARF19, key members of the transcription factor family responsible for activating AuxREs^26^, we proceeded to assess *HAT1* transcript levels in *arf7/arf19* double mutants under both mock- and IAA-treated conditions, contrasting them with those of WT plants. The marked reduction in *HAT1* transcript levels observed in the *arf7/arf19* mutants relative to WT in both treatment scenarios (Fig. 2e) underscores the participation of ARF7 and ARF19 in the transcriptional regulation of *HAT1* in response to MEcPP.

These findings reveal the involvement of MEcPP and auxin in regulating *HAT1* expression levels, compelling us to examine RSRE activities in WT and *ceh1* plants following IAA treatment. As expected, we observed a significant reduction in RSRE signals in IAA-treated plants compared to representative mock-treated plants (Fig. 2f-2g).

In summary, these data elucidate the molecular mechanisms underlying the MEcPP-mediated decrease in *HAT1* expression levels and the resulting increase in RSRE activity.

### MEcPP-mediated reduction of HAT1 interacting protein, TPL

Expanding upon the known protein-protein interaction between HAT1 and TPL, a transcriptional co-suppressor of gene expression^27^, coupled with MEcPP’s documented negative regulation of *HAT1* (Fig. 2), we postulated that MEcPP might similarly affect TPL. To address such a possibility, we initially reaffirmed TPL/HAT1 protein-protein interaction by structural modeling analyses (Fig. 3a), and further determined that TPL has an N-terminal TPR1-like fold, preceded by two WD40/YVTN repeats that assemble into a seven-blade propeller architecture (Extended Data Fig. 3a). These domains are recognized for their role in mediating protein-protein interactions^28^. Moreover, protein modeling analysis elucidated that the stability of the HAT1-TPL structure primarily relies on salt bridges formed between polar residues, a feature extensively conserved across land plants (Fig. 3a and Extended Data Fig. 3b-4c).

**Figure 3.**
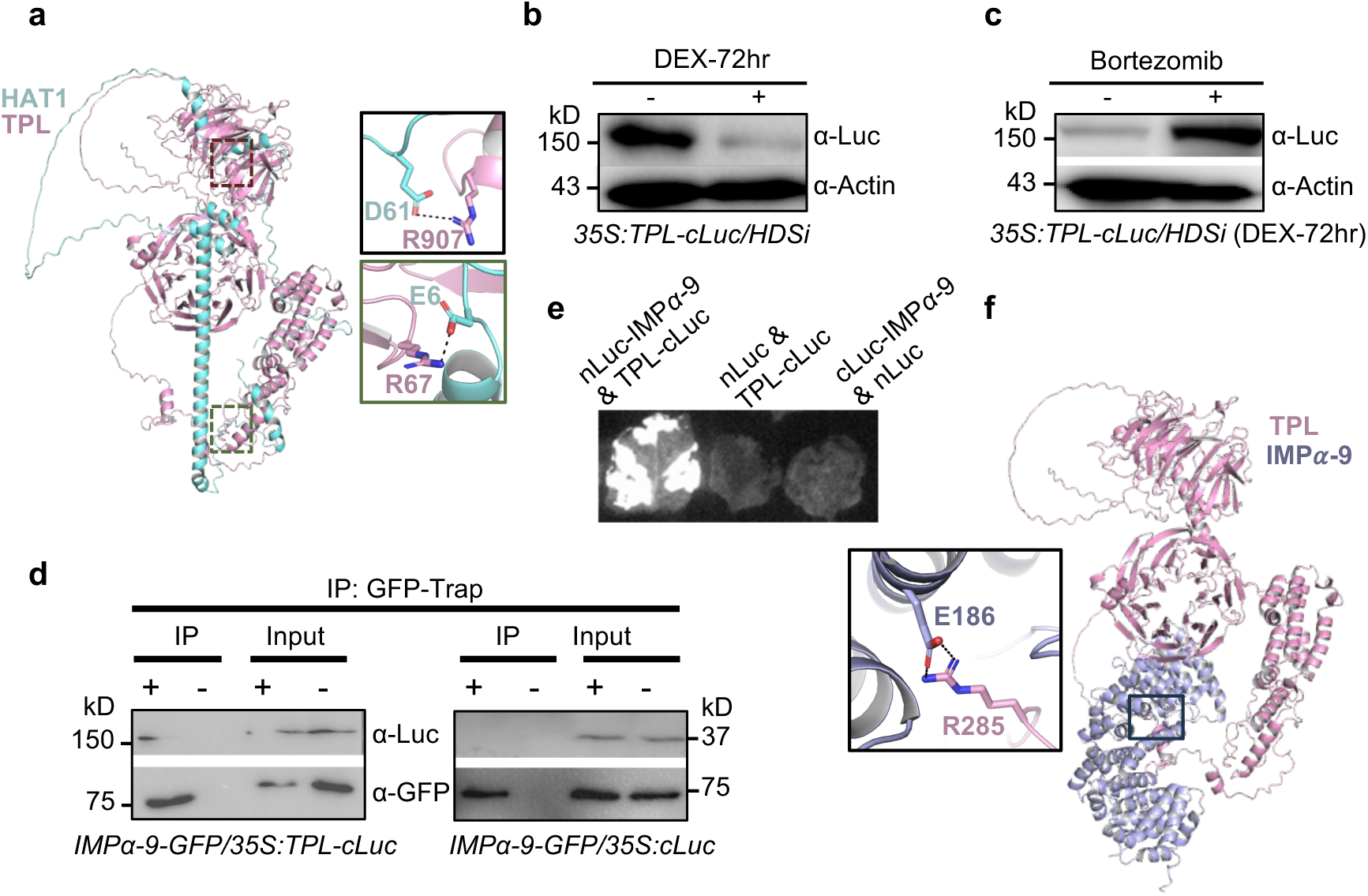
MEcPP-Mediated Reduction of HAT1-Interacting Protein TPL via the 26S Proteasome and Physical Association of TPL with IMPα-9. **a,** Highest model probability predicted by AlphaFold 3 structure of HAT1 (cyan) interaction with TPL (light pink) (left panel). Zoom-in view of the salt bridge stabilizing the HAT1-TPL complex (right panel). **b,** Protein abundance of TPL is reduced in inducible *HDSi* line post 72 h of DEX treatment (+) compared to the untreated control plants (-). **c,** Immunoblot analyses of protein abundance of TPL in 35S/HDSi plants post 72 h of DEX treatment in the absence (-) (mock, 0.1% DMSO) and presence (+) of the 26S proteasome inhibitor, bortezomib (40 μM) for 18 h, using α-Luc antibody to detect TPL-cLuc protein, and α-Actin antibody served as the loading control. **d,** The *in vivo* interaction of IMPα-9 with TPL determined by co-immunoprecipitation assay. Protein samples obtained from IMPα-9-GFP/35S were immunoprecipitated (IP) using GFP (+) and empty (-) magnetic beads followed by immunoblots performed using α-Luc. Each blot shows protein inputs before (input, right panels) and after (IP, left panels) immunoprecipitation. **e,** Split luciferase complementation assays in *Nicotiana benthamiana* leaves expressing 35S: IMPα-9 (N-terminal Luc fragment fused with IMPα- 9) and 35S (C-terminal Luc fragment fused with TPL), demonstrating the interaction between IMPα-9 and TPL. Negative controls include 35S:IMPα-9 & 35S;cLUC. **f,** Highest model probability predicted by AlphaFold 3 structure of TPL-IMPα-9 complex (right panel). Zoom-in view of the salt bridge between TPL and IMPα-9 (left panel).

Next, we examined the transcript levels of *TPL* in two-week-old WT and *ceh1* plants and found no significant differences between the two lines (Expanded Data Fig. 4). This suggests that MEcPP does not exert transcriptional regulation over TPL.

**Figure 4.**
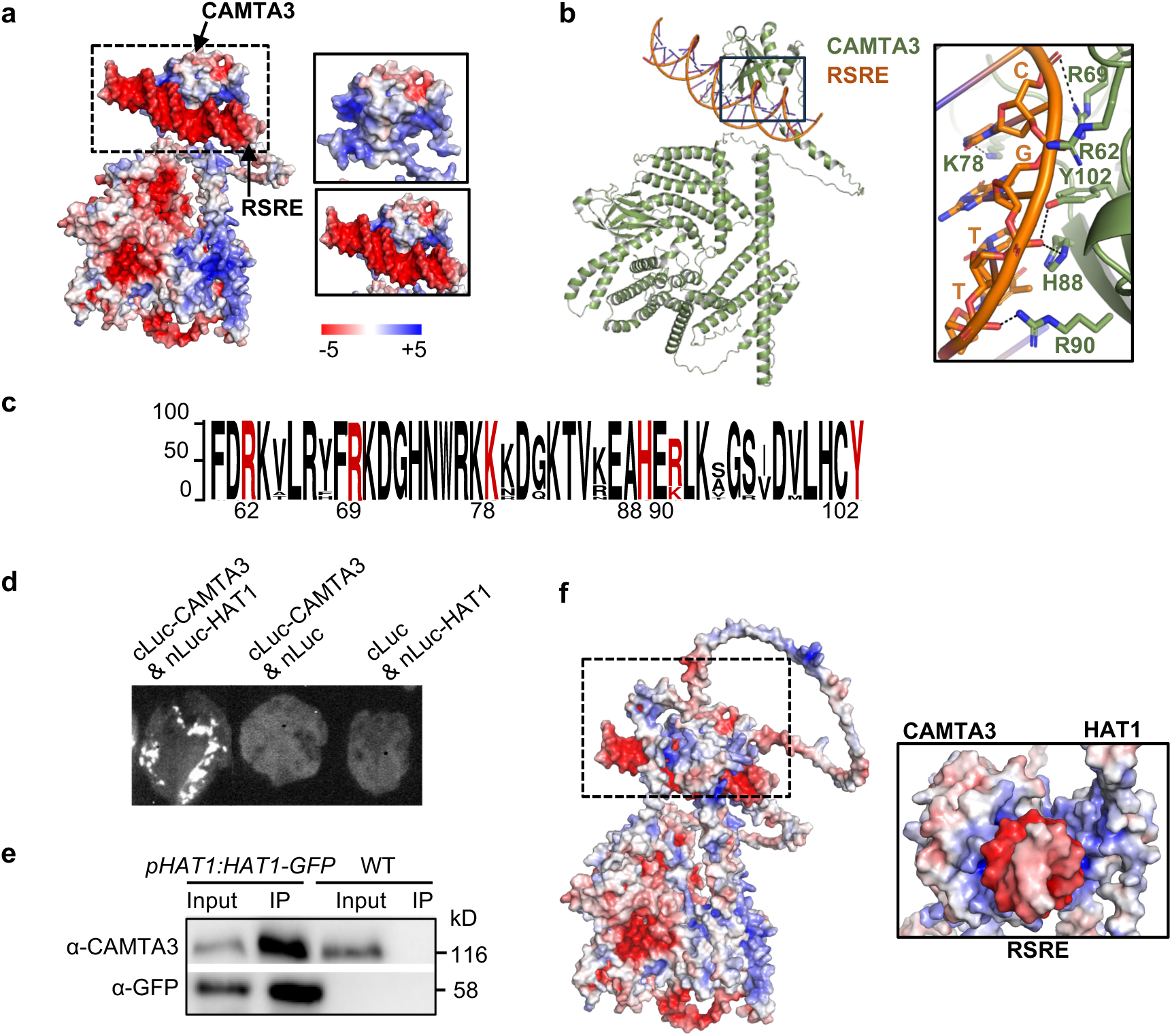
HAT1 physically interacts with the activator of RSRE, CAMTA3. **a,** Electrostatic surface charge distribution of the predicted CAMTA3-RSRE complex (left panel), and the zoom-in view of CAMTA3 (right top panel) and CAMTA3-RSRE complex (right bottom panel). The bar below shows the surface charge distribution calculated using the APBS program in PyMOL. **b,** Predicted structure of CAMTA3 (light green)-RSRE (orange) complex and the zoom-in view of hydrogen bond interaction within the CAMTA3-RSRE complex (right panel). **c,** The sequence logo generated using the protein sequences of CAMTA3 from representative land plants, highlighting the conservation of RSRE-binding sites on CAMTA3. The red amino acids are the RSRE- binding residues. Numbers under the sequence logo indicate the amino acid positions in Arabidopsis CAMTA3. The scale reflects the percentage of conservation. **d,** The representative split luciferase complementation assays in *Nicotiana benthamiana* leaves expressing 35S: C-Luc fragment fused with CAMTA3 and 35S: HAT1 fused with N-Luc fragment, demonstrating the interaction between HAT1 and CAMTA3. Negative controls include 35S:cLuc-CAMTA3 & 35S:nLuc and 35S:nLuc-HAT1 & 35S:cLuc. **e,** The *in vivo* interaction of IMPα-9 with TPL determined by co-immunoprecipitation assay. Protein samples obtained from pHAT1 and WT (Col-0) were immunoprecipitated using GFP (+) magnetic beads. Immunoblots were analyzed using α-CAMTA3. Each blot shows protein inputs before (input, left panels) and after (IP, right panels) immunoprecipitation. **f,** Predicted structure of CAMTA3 (light purple)-HAT1 (blue) and RSRE (red) complex and the zoom-in view of hydrogen bonding interaction complex (right panel).

To investigate whether MEcPP regulates TPL’s protein level, and in the absence of an available antibody, we introduced the *35S:TPL-cLuc* vector into previously generated DEX-inducible hydroxymethyl butenyl diphosphate synthase (HDS) RNAi (*HDSi*) plants. These plants accumulate MEcPP at levels similar to those observed in *ceh1* mutant after 72 h of DEX induction^29^. We subjected two-week-old *35S:TPL-cLuc/HDSi* seedlings to DEX or control mock, followed by immunoblot analyses conducted 72 h later to evaluate total TPL protein levels. The data showed a significant reduction in TPL total protein levels as evident in the DEX-induced *35S:TPL-cLuc/HDSi* seedlings compared to the mock-treated counterparts (Fig. 3b). This finding clearly establishes a link between MEcPP accumulation and the decreased TPL protein levels in DEX-induced *HDSi* seedlings.

To deepen our understanding of how MEcPP influences TPL protein abundance, we investigated the impact of bortezomib, a potent 26S proteasome inhibitor, on TPL protein levels. Two-week-old *35S:TPL-cLuc/HDSi* seedlings at 72 h after DEX treatment, were treated with either a bortezomib solution (40 μM in 0.1% DMSO) or a mock solution (0.1% DMSO) for 18 hs. Subsequent immunoblot analyses to evaluate TPL protein levels revealed a significant increase in protein abundance in the bortezomib-treated seedlings compared to their mock-treated counterparts (Fig. 3c and Extended Data Fig. 5a).

This bortezomib-heightened stability of TPL protein strengthens the notion that proteasomal degradation plays a pivotal role in the MEcPP-mediated regulation of TPL protein levels.

### TPL interacts with nuclear importin IMPα-9

In our prior investigation using genetic mutagenesis analysis, we identified nuclear importin IMPα-9 as a key suppressor of the MEcPP-mediated retrograde signal transduction pathway. Additionally, we found that MEcPP negatively affects IMPα-9 protein levels^22^. Through immunoprecipitation-mass spectrometry (IP-MS) and yeast two-hybrid library screening assays, we discovered various members of the TPL family of transcriptional co-repressors in the interactome of IMPα-9, including TPL itself^22^.

We subsequently utilized additional methods to confirm the protein-protein interaction between TPL and IMPα-9. First, we generated *pIMP-α9:IMPα-9-GFP/35S:TPL-cLuc* (referred to as *IMPα-9-GFP/35S:TPL-cLuc*) and control *pIMP-α9:IMPα-9- GFP/35S:cLuc* (referred to as *IMPα-9-GFP/35S:cLuc*) transgenic plants to verify the interaction through targeted co-immunoprecipitation (Co-IP) experiments. We used GFP-Trap and empty magnetic beads for IP of *IMPα-9-GFP* in both lines, followed by immunoblot analyses using GFP and luciferase antibodies. The clear presence of the TPL-cLuc band in the IP fraction of the *IMPα-9-GFP/35S:TPL-cLuc* line, but not in the control empty beads or the *IMPα-9-GFP/35S:cLuc* line, verified the *in vivo* interaction between TPL and IMPα-9 proteins (Fig. 3d and Extended Data Fig. 5b).

We further confirmed the interaction by luciferase reconstitution assay. We constructed vectors that fused the IMPα-9 coding sequence (CDS) with the amino-terminal fragment of the luciferase gene under the control of the 35S promoter (*35S:nLuc-IMPα-9*) and the TPL CDS with the carboxyl-terminal fragment of luciferase gene under the 35S promoter (*35S:TPL-cLuc*). Infiltrating tobacco leaves with both *35S:nLuc-IMPα-9* and *35S:TPL- cLuc* constructs resulted in the reconstitution of luciferase activity, while controls showed no activity (Fig. 3e). Lastly our structural modeling analyses validated the interaction between TPL and IMPα-9, revealing that this interaction is predominantly reinforced by the salt bridge formed between TPL^R285^ and IMPα-9^E186^, thereby stabilizing the TPL- IMPα-9 complex (Fig. 3f). Notably, these interaction sites were found to be highly conserved across land plants (Extended Data Fig. 6).

In summary, these results clearly establish the protein-protein interaction between TPL and IMPα-9.

### HAT1 interacts with CAMTA3, an activator of MEcPP-mediated RSRE induction

It is widely acknowledged that CAMTA3, activated by stress-induced MEcPP, acts as the primary activator of RSRE^2,30^. The predicted structure of CAMAT3 reveals conserved functional domains such as N-terminal DNA binding domain, tandem ankyrin repeats that are involved in protein-protein interaction, IQ motifs which are implicated in the association of Ca^2+^ loaded CaM and CaM-like proteins (CML) to CAMTA^31^ (Extended Data 7a). Detailed structural analysis further elucidated the hydrogen bonding interactions between CAMTA3 and RSRE (Fig. 4a-4b). Interestingly, evolutionary analyses suggest that these interaction sites, some of which have been confirmed through point mutations^30^, are highly conserved across land plants (Fig. 4c and Extended Data 7b).

Given MEcPP’s established role in CAMTA3 activation, we explored whether the RSRE suppressor, HAT1, interacts with CAMTA3 under normal conditions or if they operate independently. To test these possibilities, we initially utilized a luciferase reconstitution assay and constructed vectors expressing fusion proteins: the *HAT1* coding sequence with the amino-terminal fragment of the luciferase gene (*nLuc-HAT1*) under the 35S promoter, and the CAMTA3 with the carboxyl-terminal fragment of luciferase (*cLuc-CAMTA3*) under the 35S promoter. Co-infiltration of tobacco leaves with these constructs showed the reconstitution of luciferase activity exclusively in leaves infiltrated with both *nLuc- HAT1* and *cLuc-CAMTA3*, supporting their interaction (Fig. 4d).

To further confirm this interaction, we generated transgenic Arabidopsis plants expressing HAT1-GFP under its native promoter (*pHAT1:HAT1-GFP*) and conducted Co-IP experiments. Utilizing GFP-Trap magnetic beads, we isolated immunoprecipitation fractions from the transgenic and wild-type control lines, followed by immunoblot analyses with antibodies against GFP and CAMTA3. The detection of CAMTA3 in the immunoprecipitate from *pHAT1:HAT1-GFP* plants, but not from wild-type plants, provided additional evidence for the *in vivo* interaction between HAT1 and CAMTA3 (Fig. 4e). Additionally, structural analysis suggests that RSRE is positioned between HAT1 and CAMTA3. HAT1, together with the positively charged DNA-binding domain of CAMTA3 (Extended Data Fig. 7a), engages in electrostatic interactions with RSRE (Fig. 4f). Evolutionary analyses spanning land plants also unveil the conservation of interaction sites among HAT1-RSRE, CAMTA3-RSRE, and CAMTA3-HAT1 (Extended Data Figs. 2, 7b, and 8).

In conclusion, these findings definitively establish the protein-protein interaction between HAT1 and CAMTA3, indicating that HAT1 binds to and hinders CAMTA3’s access to RSRE motifs in the promoter region of stress-response genes under normal conditions.

### MEcPP potentiates extracellular Ca^2+^ influx

We have established the pivotal role of MEcPP in activating RSRE, which was subsequently nullified by the presence of the Ca^2+^ chelator EGTA^2^. This highlights the indispensable function of Ca^2+^ for CAMTA3 activity, leading to the notion of MEcPP as a regulator for Ca^2+^ release, thereby facilitating CAMTA3 activation^2,32^. To investigate this notion, we utilized our previously reported multi-compartmental Ca^2+^ sensor construct (CamelliA_NucG/PmG/CytR) to simultaneously image Ca^2+^ dynamics in the nucleus, sub-plasma membrane (sub-PM), and cytosol^33^. We expressed this sensor in both WT plants and high MEcPP-accumulating *ceh1* mutant and examined the Ca^2+^ dynamics in both backgrounds. The results showed significantly higher and more frequent Ca^2+^ fluctuations in the nucleus and cytosol of epidermal cells of the *ceh1* mutant compared to WT plants (Fig. 5a, and Extended Data Movie 1-2), suggesting contribution of elevated MEcPP levels to much stronger Ca^2+^ fluctuations in the *ceh1* mutant.

**Figure 5.**
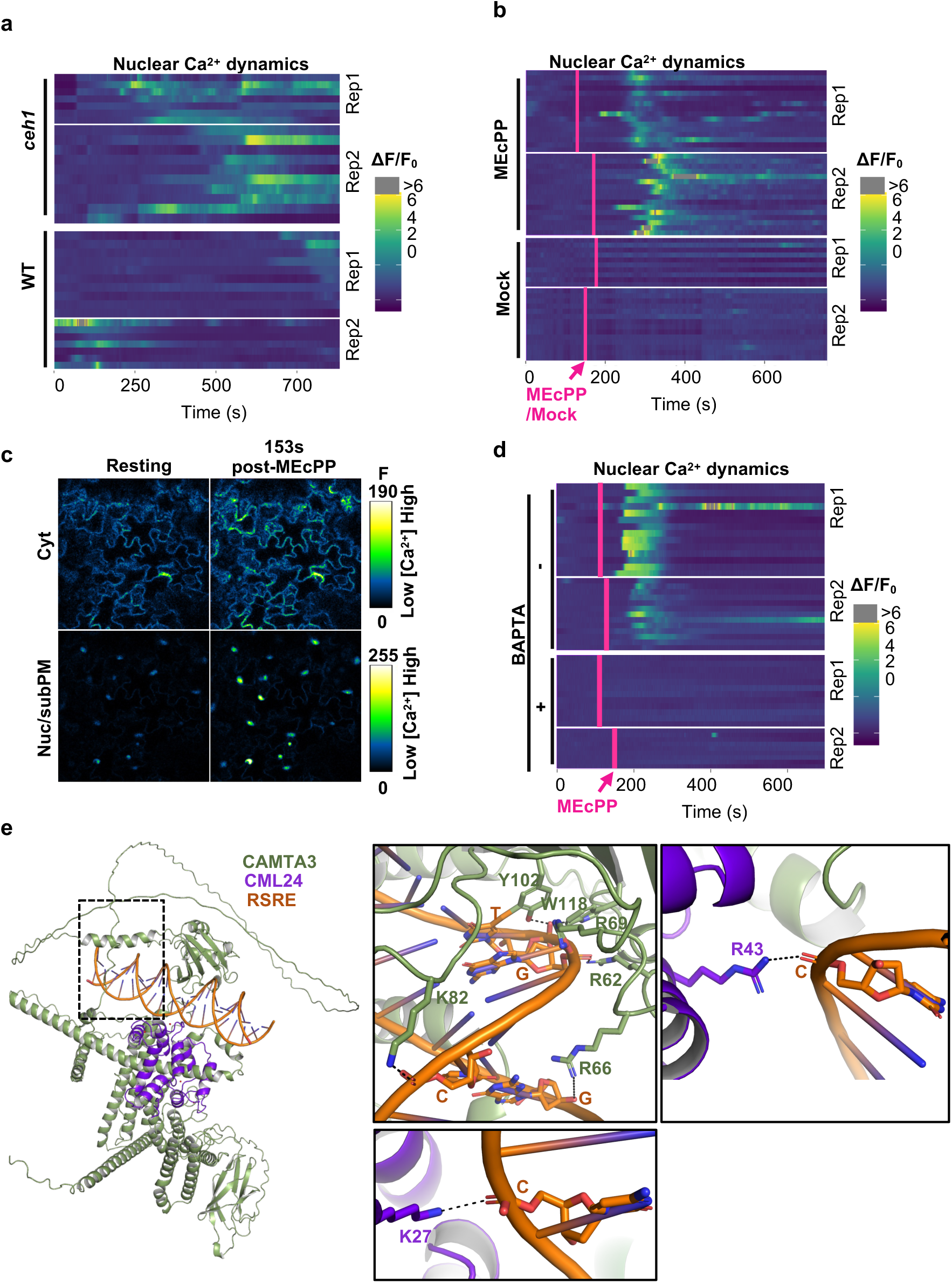
MEcPP triggers influx of extracellular Ca²⁺ to activate CAMTA3. **a-b,** Heatmaps show the induction of a nuclear Ca²⁺ signature in wild type (WT) and MEcPP-accumulating mutant *ceh1* (**a**), and in seedlings treated exogenously with MEcPP or a mock treatment (**b**). The heatmap traces the relative changes in fluorescence intensity in the nuclei of the triple Ca²⁺ sensor CamelliA_NucG/PmG/CytR in WT and the constitutively MEcPP-producing line (*ceh1*), and WT before and after MEcPP/mock treatment (**b**), with the red line indicating the time of MEcPP and Mock application. **c,** Representative images show MEcPP-triggered Ca²⁺ dynamics in the cytosol and nucleus/sub-PM (plasma membrane) of WT prior (Resting) and post MEcPP treatment. **d,** Induction of the nuclear Ca²⁺ signature by exogenous application of MEcPP in the presence or absence of 10 mM BAPTA. (**e**) Interaction of CAMTA3 and CML24 with RSRE (left panel). Zoom-in view of the interacting amino acid residues of CAMTA3 and CML24 with RSRE (right panel). Dashed lines indicate the hydrogen bond interactions. Each line in the heatmaps represents the time trace of Ca²⁺ dynamics in an individual nucleus, with two representative heatmaps of time traces (Rep1 and Rep2) shown. Scale bar is 20 µm in (**c**).

To investigate the specificity of MEcPP in potentiating these Ca^2+^ signatures, we examined the Ca^2+^ signature in WT plants treated with either a mock or exogenous MEcPP. The results demonstrated that exogenously applied MEcPP, but not the mock treatment, induced both nuclear and cytosolic Ca^2+^ elevation in epidermal cells of true leaves (Fig. 5b-c and Extended Data Movie 3-4).

Furthermore, we employed the commonly used calcium chelator, BAPTA (1,2-bis(o-aminophenoxy) ethane-N,N,N’,N’-tetraacetic acid), which sequesters Ca^2+^ by forming stable complexes through its multiple oxygen atoms, effectively reducing the concentration of free Ca^2+^ in the cellular environment. Our data unequivocally demonstrate the abolition of Ca^2+^ influx in the presence of BAPTA (Fig. 5d and Extended Data Movie 5).

Given the constitutive expression of RSRE in the *ceh1* mutant background and the significant increase in Calmodulin-like protein 24 (CML24) expression levels in *ceh1*^1,2^, along with the role of CML24 as a potential Ca^2+^ sensor and the function of Ca^2+^- Calmodulin in activating CAMTA3, we were prompted to further analyze the structure of Ca^2+^-loaded CML24 in complex with CAMTA3 (Fig. 5e and Extended Data Fig. 9a). These analyses clearly revealed the four conserved EF-hand motifs of CML24 responsible for Ca^2+^ binding^31^, primarily stabilized by aspartic/glutamic acid and asparagine residues (Extended Data Fig. 9b). Furthermore, the structure of CML24 features a dumbbell-shaped architecture with N- and C-terminal domains connected by a flexible linker that wraps around CAMTA3, leading to the binding of CAMTA3’s IQ-motif peptide at the center of the N- and C-terminal domains. This interaction is primarily stabilized by polar salt bridge interactions (Extended Data Fig. 9c). These conformational changes, unlike those in the repressor-bound CAMTA3 complex (Fig. 4f and Extended Data Fig. 10), result in the activation of CAMTA3, allowing it to fully access its binding site, RSRE (Fig. 5e), thereby prompting the expression of stress-induced genes. Collectively, these findings further support our previous notion that MEcPP potentiates extracellular Ca^2+^ influx, elevating cytosolic and nuclear Ca^2+^ levels. This, in turn, activates Ca^2+^-CML-dependent activation of CAMTA3, leading to the transcription of stress response genes.

## Discussion

Understanding the regulatory components of the General Stress Response (GSR), also known as the cellular stress response or core stress response, is essential for deciphering the fundamental principles governing early stress response mechanisms impacting cellular homeostasis, providing cross-protection, where exposure to one stressor can confer resistance to another, and enabling tailored adaptive responses to specific stressors, among other functions. Extensive research across bacteria, fungi, animals, and plants has confirmed the evolutionary conservation of GSR molecular components across all domains of life, highlighting their critical role in survival under stressful conditions^23,34,35^.

The rapid activation of the GSR, often occurring within minutes of exposure to stimuli, demands efficient perception and integration of signal transduction, alteration of pre-existing transcription factors, and epigenetic modifications. While specific regulatory elements may vary among organisms, the underlying principles of stress detection, signal transduction, and adaptive responses remain evolutionarily conserved. Moreover, within the same organism, shared regulatory infrastructure allows for effective management and response to diverse environmental stressors. This conservation has spurred extensive inquiries into the mechanisms involved, providing deeper insights into how organisms maintain cellular homeostasis and adapt to environmental challenges.

In this study, we investigated the intricate mechanisms governing the transmission of stress-induced plastidial retrograde metabolite MEcPP signaling to the nucleus, with a particular focus on its crucial role in initiating the expression of GSR genes, notably through activating the GSR transcriptional hub RSRE. Our investigation illuminates MEcPP’s dynamic roles in coordinating master regulators of adaptive responses through two distinct pathways. MEcPP disrupts the HAT1/TPL/IMPα-9 suppressor complex through a dual mechanism. Firstly, it reduces levels of the hormone IAA, thereby diminishing IAA-dependent expression of HAT1. Simultaneously, MEcPP promotes TPL/IMPα-9 degradation via the 26S proteasome machinery. Additionally, our recent discovery of MEcPP-induced ASK1 protein accumulation, a crucial component in the SKP1-Cullin 1-F-box (SCF) E3 ubiquitin ligase complex, unveils another facet of MEcPP’s regulatory role^22^. ASK1 facilitates IMPα-9 degradation, further disintegrating the HAT1/TPL/IMPα-9 suppressor complex. This releases the sequestered activator CAMAT3, enabling it to bind to RSRE and initiate stress-responsive gene expression. Indeed, the predicted structure model of the RSRE-CAMTA3-HAT1-TPL-IMPα-9 complex reinforces our experimental data (Extended Data Fig. 10).

Remarkably, this mechanism shares similarities with established signaling pathways such as jasmonate (JA) and abscisic acid (ABA) pathways. In JA signaling, JAZ (Jasmonate ZIM-Domain) proteins act as repressors inhibiting transcription factors like MYC2 responsible for defense gene expression. However, during stress, increased JA levels trigger JAZ degradation through the Skp/Cullin/F-box complex (SCF^COI^^1^)-dependent 26S proteasome pathway, freeing MYC2 to activate defense-related genes^36,37^. Similarly, in the ABA signaling pathway, PP2C (Protein Phosphatase 2C) proteins dephosphorylate and repress SnRK2 kinases, but under stress ABA inhibits PP2C, allowing SnRK2 kinases to activate downstream targets and initiate ABA-responsive gene expression^38^.

These signaling pathways collectively illustrate a coordinated regulation amounting to relieving repression and activating key components to mount an effective response to stress.

Besides its role in releasing transcription factors and alleviating repression, MEcPP also enhances the influx of Ca^2+^, a crucial step for activating CAMTA3. This adds complexity to the regulatory network of retrograde signaling in adaptive responses. This discovery raises an intriguing question about the nature of calcium channels and the mechanism by which MEcPP activates them. Unraveling these mechanisms provides insights into how plants integrate various signals to orchestrate effective responses to environmental stresses.

In summary this report outlines the intricate network of transcriptional activities that balance growth and stress responses, illustrated in the simplified schematics (Fig. 6). Under unstressed conditions, the HAT1 complex binds to the RSRE motif present in 30% of stress response genes, repressing their expression. However, stress-induced accumulation of MEcPP triggers a synchronized dual-channel mechanism that removes the repressor complex while activating the activator (CAMTA3) by enhancing nuclear calcium influx. This interplay among retrograde signaling, nuclear transport, transcription regulation, and cellular ion signaling underscores the pivotal and multifaceted roles of the signaling metabolite. Our findings significantly advance the understanding of retrograde metabolite signaling in coordinating plastid-to-nucleus communication, emphasizing the central role of plastids as regulatory hubs in orchestrating plant responses to stress.

**Figure 6.**
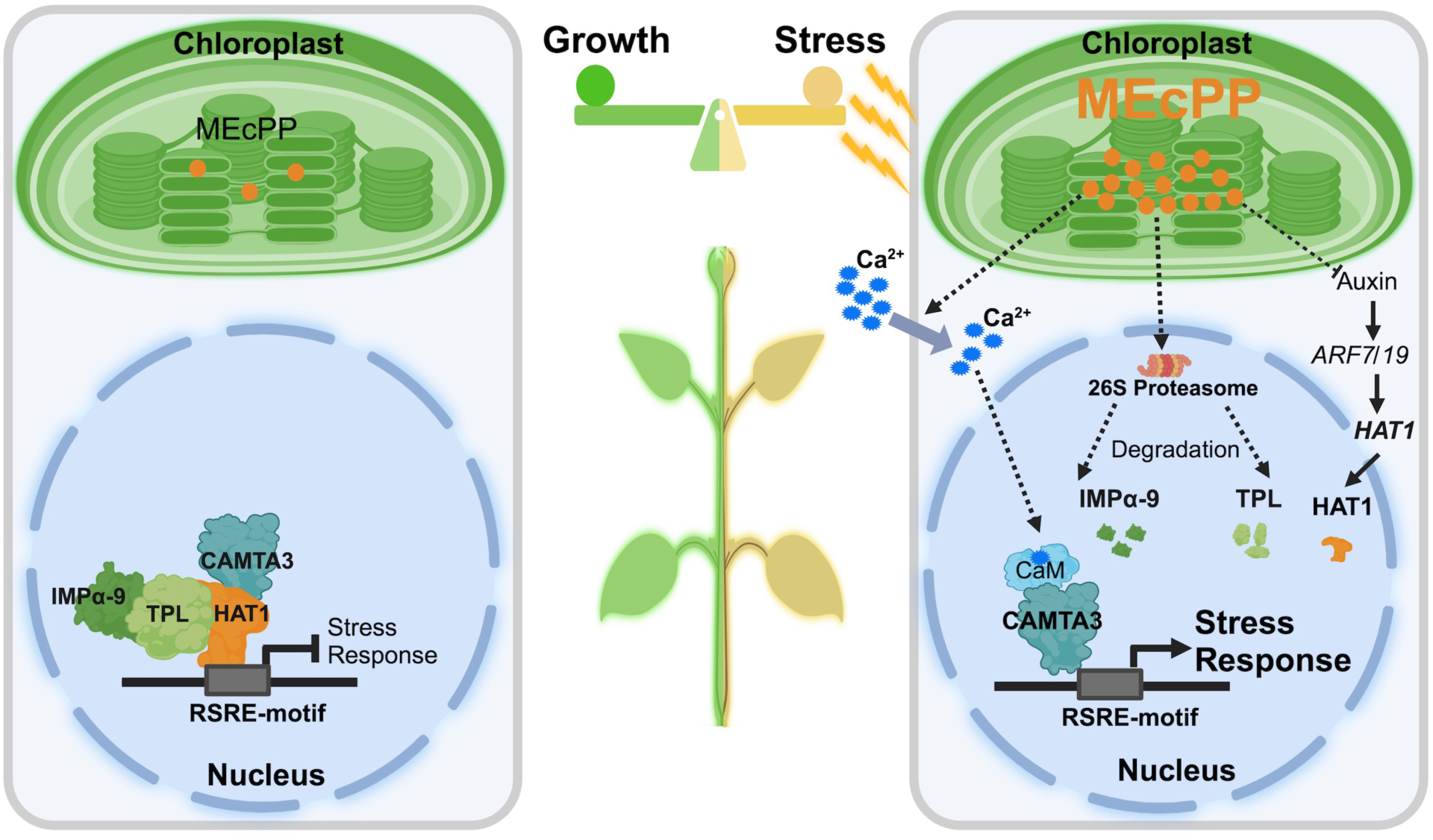
Plastidial MEcPP: A Gatekeeper of Growth and Stress Response Balance in Plants. Plastids play a crucial role in balancing growth and stress responses through the retrograde signaling metabolite MEcPP. Under standard conditions, a repressor complex composed of HAT1, TPL, and IMPα-9 binds to the transcriptional stress hub RSRE, sequestering the activator CAMTA3 and repressing stress response genes. When stress occurs, the accumulation of MEcPP triggers the dismantling of the repressor complex, releasing CAMTA3 and activating it through calmodulin (CaM) binding facilitated by increased nuclear calcium influx from extracellular space. This synchronized dual- channel mechanism underscores the pivotal role of plastids and MEcPP in regulating plant stress responses.

## Materials and Methods

### Plant material and growth condition

All *Arabidopsis thaliana* mutants and transgenic lines are in the Columbia (*Col-0*) ecotype. Plants were grown in 16-h light/8-h dark cycles at ∼22 ^°^C. Two-week-old seedlings grown on ½ MS plates were used for all the experiments. The *ceh1*, HDSi lines were developed previously^1,29^. OX-*ARF* seeds were obtained from Professor Lucia C. Strader at Washington University. Transgenic *pIMPα-9:IMPα-9-GFP*/*35S:TPL-cLuc*, *pIMPα-9:IMPα-9-GFP*/*35S:cLuc*, *35S:HAT1*, CRISPR-*hat1*, *35S:TPL-cLuc/HDSi* and *pHAT1:HAT1-GFP* lines were generated in this study. Two-week-old seedlings were treated with IAA (10 μM) for 30 minutes to extract RNA, or 1 h for luciferase signals measurements. Forceps were used to mechanically wound leaves. The RSRE::LUC signals were checked by using the CCD camera as previously described^39^.

### Accession Numbers

The gene sequences mentioned in this article are available in the Arabidopsis Genome Initiative Information Resource (http://www.arabidopsis.org/). The locus numbers corresponding to the genes are as follows: *IMPα-9* (AT5G03070), *TPL* (AT1G15750), *HAT1* (AT4G17460), *HDS* (AT5G60600) and *CAMTA3* (AT2G22300).

### Yeast one-hybridization assay

The Gold Yeast One-Hybrid Library Screening System (#630491; Takara, Shiga, Japan) was used for screening RSRE motif binding proteins. The sequence of RSRE motif (the ‘DNA bait’) was cloned to create the 4xRSRE::LacZ and 4xRSRE::Luciferase reporter constructs. X-gal was used to detect LacZ activity. The Andor Technology CCD camera was used to detect luciferase activity signals.

### MEcPP extraction and analyses

MEcPP were extracted and analyzed as previously described^40^.

### Quantification of gene expression

Total RNA was extracted from two-week-old plants using the Aurum Total RNA Mini Kit (Bio-Rad). Reverse transcription of 1 μg of total RNA was carried out using the iScript Reverse Transcription Supermix for RT-PCR according to the manufacturer’s instructions (Bio-Rad). The resulting complementary DNA (cDNA) was used as the template for subsequent analysis. Real-time PCR was performed using the SsoAdvanced Universal SYBR Green Supermix on a CFX96 real-time PCR detection system (Bio-Rad). The reference gene *AT4G26410* was used as an internal control.

### Statistical analysis

Two-tailed Student’s *t-*test and ANOVA test with Tukey’s post hoc test were used for the statistical analyses, the asterisks and different letters on figures denote statistical significance (*P* ≤ 0.05).

### Bortezomib treatment

Two-week-old *pIMPα-9:IMPα-9-GFP/HDSi* and *pIMPα-9:impα-9-GFP/HDSi* seedlings 72 h post DEX treatment were used for these analyses. Leaves were cut as small pieces and submerged in bortezomib (40 μM, dissolved in 0.1% DMSO) or in mock (0.1% DMSO) buffer for 18 h on a shaker at room temperature.

### Agro-infiltration-based transient assays in *Nicotiana benthamiana*

The *Nicotiana benthamiana* transient assay was employed to investigate the RSRE activity and protein-protein interactions. Vectors pENTR/D-TOPO (Invitrogen) and Gateway systems were used for vector construction. The vectors carrying C/N-terminal luciferase fused with *IMPα-9*, *TPL*, *HAT1* and *CAMTA3*, and luciferase fused with RSRE motif were introduced into *Agrobacterium* GV3101, which were then used for leaf infiltration in *N. benthamiana*. After 48- to 72 h of infiltration, tobacco leaves were sprayed with 1 mM luciferin solution. The Andor Technology CCD camera was used to detect luciferase activity signals. Images were acquired every 5 min for 3 h for tobacco plants. The Andor Technology software was used to quantify the luciferase activity.

### Protein extraction and immuno-blot analyses

Two-week-old seedlings were collected and ground in liquid nitrogen and suspended in 2x SDS lysis buffer and boiled (100 ^°^C, 10 min) for total protein extraction. Proteins were then separated on 10% SDS-PAGE gel and transferred onto the nitrocellulose membrane. The monoclonal α-GFP (1:1000, Roche), the monoclonal α-Luciferase (1:1000, Sigma), α-CAMTA3 (1:500) antibodies^41^ were used to detect the respectively tagged proteins. And the monoclonal α-Actin (1:10,000, Sigma) antibody was used as the loading control. The goat anti-mouse (1:3000) IgG (H+L)-HRP (1:3000) and goat anti-rabbit IgG (H+L)- HRP (1:3000) conjugated secondary antibodies (Bio-Rad) were used.

### Co-Immunoprecipitation (Co-IP)

Two-week-old plants were grounded and suspended in extraction buffer (50 mM Tris-HCl at pH 7.5, 150 mM NaCl, 10% glycerol, 0.1% NP-40 and protease inhibitor cocktail) at 4 ^°^C for 30 min. The protein suspensions were then centrifuged at 12,000*g* for 10 min at 4 ^°^C, and the precipitation were then filtered out using a 100 μm Nylon Mesh. The supernatant was incubated with GFP-Trap or empty magnetic (Chromotek) beads for 2 h at 4 ^°^C. Beads were then collected by using a magnetic column and washed five times with 2x extraction buffer.

For Co-IP experiments, proteins were eluted from GFP-trap beads using 2x SDS lysis buffer (50 mM Tris-HCl at pH 6.8, 2% SDS, 10% glycerol, 0.1% bromophenol blue, 1% 2-mercaptoethanol) and boiled at 100 ^°^C for 10 minutes. Proteins were then separated on a 10% SDS-PAGE gel and subsequently transferred (20% methanol, 25 mM Tris, 192 mM glycine, pH 8.3) onto nitrocellulose membrane for probing with the corresponding antibodies.

### Ca^2+^ dynamics imaging in epidermal cells

To image the Ca^2+^ dynamics in *ceh1* mutant plants, we first introduced the triple Ca^2+^ sensor line CamelliA_NucG/PmG/CytR expressing nuclear- and PM-targeted Ca^2+^ sensor GCaMP6f and cytosolic-targeted Ca^2+^ sensor jRGECO1a^33^ into *ceh1* mutant background by crossing. Seeds of the triple Ca^2+^ sensor line in WT and *ceh1* background were germinated on 0.5 × Murashige and Skoog (MS) solid medium without sucrose and buffered with 0.05% MES to pH5.8. Cotyledons from 6 day-post-stratification seedlings were mounted onto glass-bottomed chamber slide using Hollister medical adhesive and incubated in leaf imaging buffer (5 mM KCl, 50 µM CaCl_2_, 10 mM MES pH6.15) in an illuminated Percival growth chamber for 2 hours. Subsequently, the cotyledon samples were imaged under an upright Zeiss confocal microscope LSM 880 equipped with a Zeiss W Plan-Apochromat 40x/1.0 DIC water dipping objective lens in a “top-imaging” setup as described previously^33^. Green-colored Ca^2+^ sensors (CamelliA_NucG/PmG) are excited with a 488 nm laser line and the emission is collected between 493-550 nm; red-colored Ca^2+^ sensor (CamelliA_CytR) is excited with a 561 nm laser line and its emission is collected between 566-635 nm. Image resolution is set to 512 x 512 pixel and scan speed 0.94556 second per frame.

To monitor Ca^2+^ dynamics in WT plant treated with exogenous applied MEcPP, true leaves of the 14 day-post-stratification seedlings of the aforementioned triple Ca^2+^ sensor plant were first attached to the glass bottom chamber slide using the medical adhesive and immediately covered with 100 µl of leaf imaging buffer (5 mM KCl, 50 µM MgCl_2_, 10 mM MES pH6.15). The mounted leaves were rested for at least 2 hours in an illuminated Percival growth chamber before imaging under the identical imaging setting as described earlier. Leaf samples were first imaged for ∼120s prior to MEcPP treatment, then were imaged for another ∼600s after adding 100 µL of 200 µM MEcPP (final concentration of 100 µM) or mock (200 µM NH_4_Cl) diluted in leaf imaging buffer (indicated by the red lines in Fig. 5b). To chelate extracellular Ca^2+^, 10 mM BAPTA is included in the imaging buffer during the initial 2-hour resting incubation before imaging.

### Analysis of Ca^2+^ imaging data

Each nucleus in the time-series images was segmented manually in the FluoroSNNAP code^42^ in MATLAB (R2024a) and mean fluorescence intensity of each nucleus is analyzed and quantified using the same tool. Baseline fluorescence intensity (F_0_) is defined as the average intensity of each nucleus during the resting period before any treatments. Relative fluorescence intensity changes over time is then calculated using the equation (F-F_0_)/F_0_ in GraphPad Prism, and is plotted as heatmap using PlotTwist (huygens.science.uva.nl/PlotTwist)^43^. Each line of the heatmap represents the relative fluorescence intensity over time of a single nucleus.

### Protein structural modeling and analysis

The predicted 3D models were generated using the Alpha-Fold 3 module with the Arabidopsis sequences for HAT1 (UniProt P46600), CAMTA3 (UniProt Q8GSA7), TPL (UniProt Q94AI7), CML24 (UniProt P25070) and IMPα9 (UniProt F4KF65). The illustration and analysis of the models were produced using PyMOL v3.

## Acknowledgements

This work was supported by National Institutes of Health (NIH) R01GM107311-8, National Science Foundation (NSF) 2104365 grants, and by Dr. John W. Leibacher and Mrs. Kathy Cookson endowed chair funds to KD. This work was also in part supported by NSF-CAREER 2047396 and 2139805 grants, and in part by the U.S. Department of Energy, Office of Science, Biological and Environmental Research, Genomic Science Program under grant number DE-SC0023158 to NS.

## Extended Data

Extended Data Figure 1. Schematic presentations of two independent *HAT1* CRISPR deletion sites in CRISPR-*hat1*-*1* and CRISPR-*hat1*-*2 lines*.

**Extended Data Figure 2. Evolutionary conservation of predicted HAT1-RSRE interaction sites in land plants.**

**Extended Data Figure 3. Similar TPL transcript levels in WT and *ceh1* plants.**

**Extended Data Figure 4. Predicted protein structure of TPL, and evolutionary conservation of predicted HAT1-TPL interaction sites in land plants.**

**Extended Data Figure 5. The original immunoblots.**

**Extended Data Figure 6. Evolutionary conservation of predicted TPL-IMP*α*-9 interaction sites in land plants.**

**Extended Data Figure 7. Predicted protein structure and conservation sites of CAMTA3.**

**Extended Data Figure 8. Evolutionary conservation of predicted CAMTA3-HAT1 interaction sites in land plants.**

**Extended Data Figure 9. Predicted protein structure of RSRE-CML24-CAMTA3 complex.**

**Extended Data Figure 10. Predicted protein structure of RSRE-CAMTA3-HAT1- TPL-IMPα-9 complex.**

**Extended Data Movie S1 Cytosolic, nuclear, and sub-PM Ca^2+^ dynamics in *ceh1* cotyledon epidermal cells.**

**Extended Data Movie S2 Cytosolic, nuclear, and sub-PM Ca^2+^ dynamics in WT cotyledon epidermal cells.**

**Extended Data Movie S3 Cytosolic, nuclear, and sub-PM Ca^2+^ dynamics in epidermal cells of true leaf treated with MEcPP.**

**Extended Data Movie S4 Cytosolic, nuclear, and sub-PM Ca^2+^ dynamics in epidermal cells of true leaf after mock treatment.**

**Extended Data Movie S5 MEcPP failed to trigger cytosolic, nuclear, and sub-PM Ca^2+^ elevation in epidermal cells of true leaf after BAPTA treatment.**

## Notes

### Competing Interest Statement

The authors have declared no competing interest.

