## Supplementary material for "Integrated Dual-Channel Retrograde Signaling Directs Stress Responses by Degrading the HAT1/TPL/IMPα-9 Suppressor Complex and Activating CAMTA3": Figures

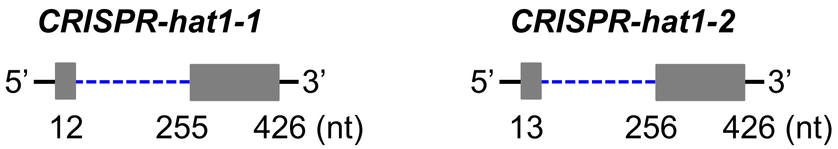


**Extended Data Figure 1. Schematic presentations of two independent *HAT1* CRISPR deletion sites in CRISPR-*hat1-1* and CRISPR-*hat1-2 lines*.**
The blue-dotted lines show the deleted region within the coding sequence of *HAT1*.

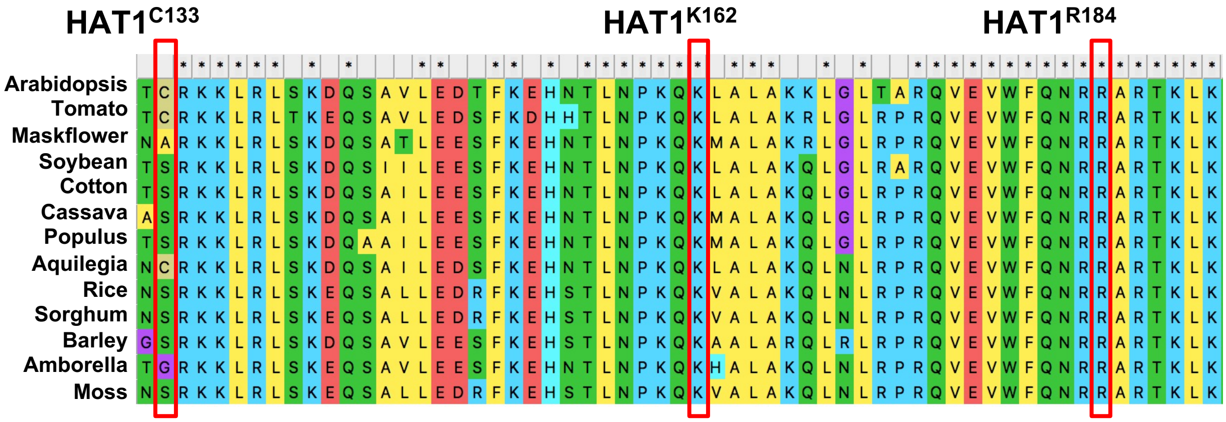


**Extended Data Figure 2. Evolutionary conservation of predicted HAT1-RSRE interaction sites in land plants.**

Protein sequencing alignments of HAT1 across representative land plant species highlights the conserved amino acids K162 and R184 sites, which are implicated in interacting with RSER.

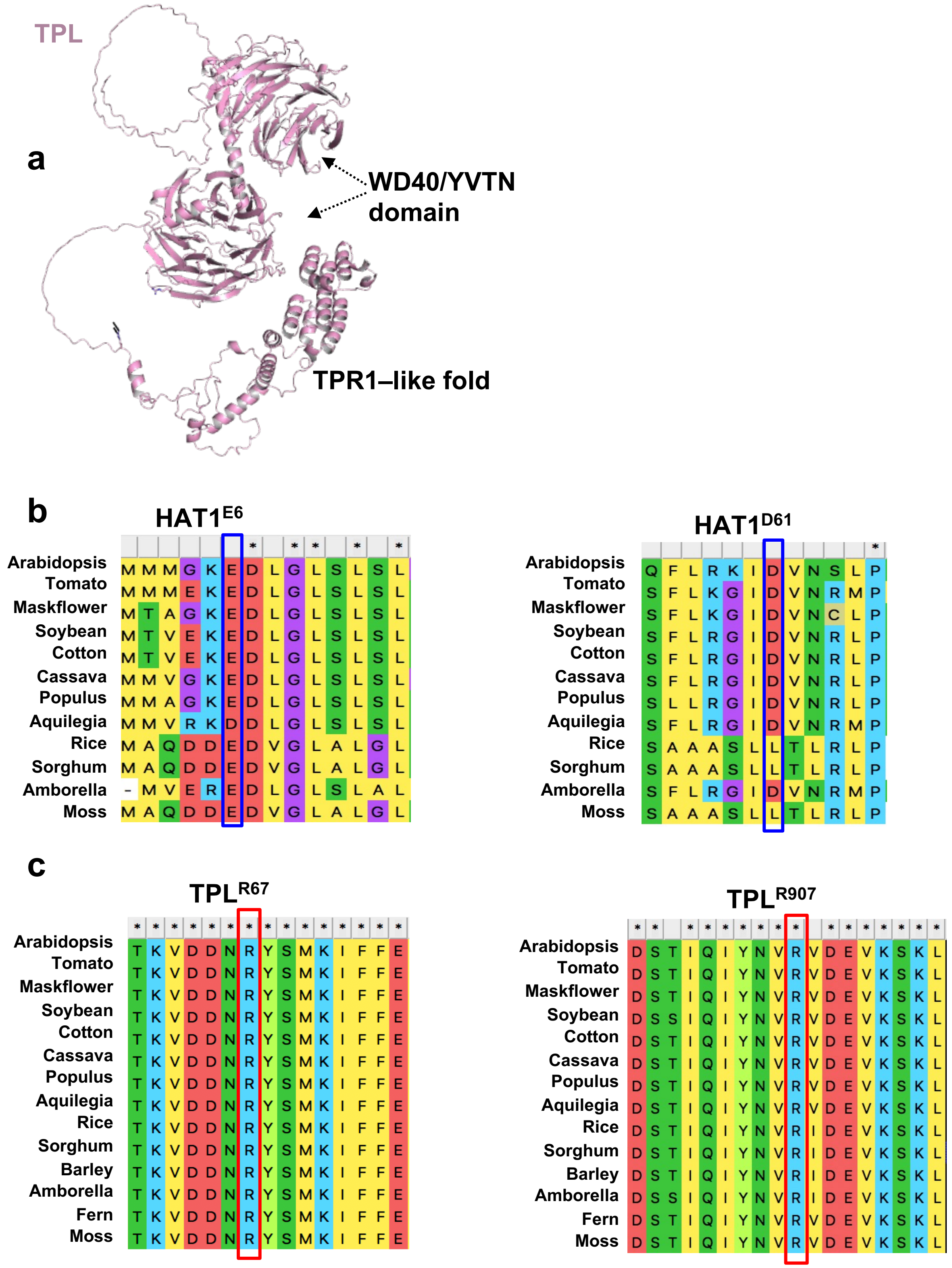


**Extended Data Figure 3. Predicted protein structure of TPL, and evolutionary conservation of predicted HAT1-TPL interaction sites in land plants.**
**a,** Predicted structure of TPL (*pink*)protein. TPL protein consist of TPR1-like fold at its N-terminus preceded by two WD40/YVTN repeat like motifs that form a seven-bladed propeller architecture. The structural analysis were based on highest model probability predicted by AlphaFold 3. **b,** Protein sequence alignments of HAT1across representative land plant species shows the conservation of the E6 and D61 site, the predicted interaction site with TPL. **c,** Protein sequence alignments of TPL across representative land plant species shows the conservation of the R67 and R907 sites, the predicted interaction site with HAT1.

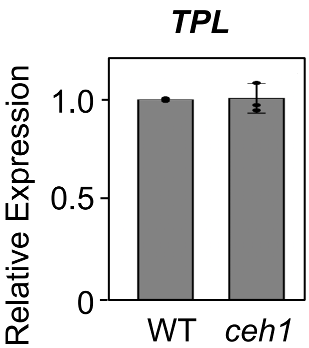


**Extended Data Figure 4. Similar TPL transcript levels in WT and *ceh1* plants.**

a

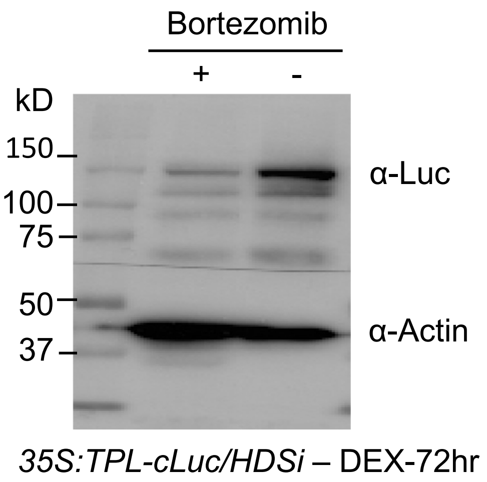


b

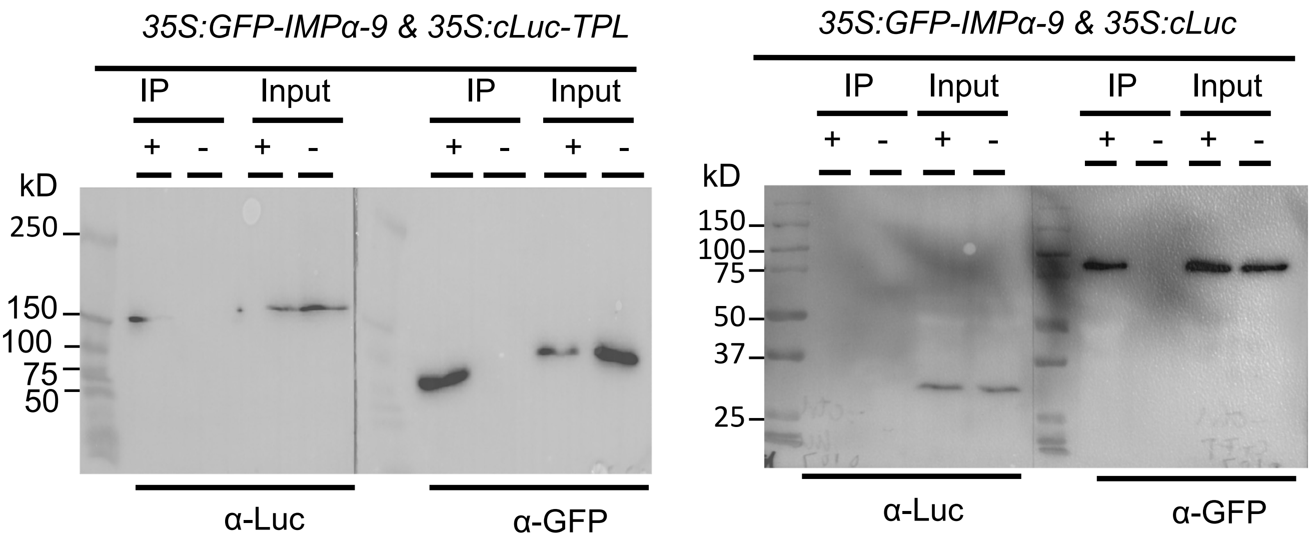


**Extended Data Figure 5. The original immunoblots.**
**a**, The original immunoblot to show that IMPα-9 physically interacts with TPL. The *in vivo* interaction of IMPα-9 with TPL determined by co-immunoprecipitation assay. Proteins extracted from tobacco leaves are immuneprecipitated using GFP (+) and empty (-) magnetic beads. Each blot shows protein inputs before (input, right panels) and after (IP, left panels) immunoprecipitation. **b**, The original immunoblot shows that bortezomib treatment stabilizes the protein abundance of TPL. Protein abundance of TPL in *35S:TPL-cLuc/HDSi* plants post 72 h of DEX treatment in the absence (-) (mock, 0.1% DMSO) and the presence (+) of the 26S proteasome inhibitor, bortezomib (40 μm) for 18 h. α-Luc antibody is used to detect TPL-cLuc protein, and α-Actin antibody served as the loading control.

**a**

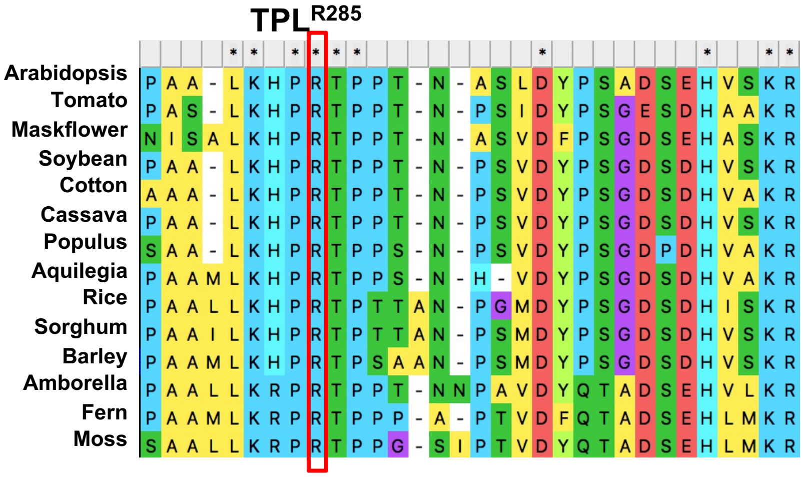


**b**

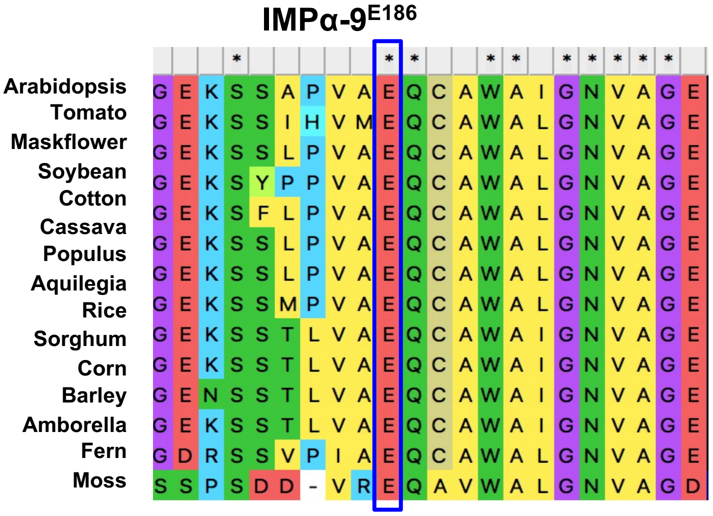


**Extended Data Figure 6. Evolutionary conservation of predicted TPL-IMPα-9 interaction sites in land plants.**

**a,** Protein sequence alignment of IMPα-9 across representative land plant species shows the conservation of the E186 site, the predicted interaction site with TPL. **b,** Protein sequence alignment of TPL across representative land plant species shows the conservation of the R285 site, the predicted interaction site with IMPα-9.

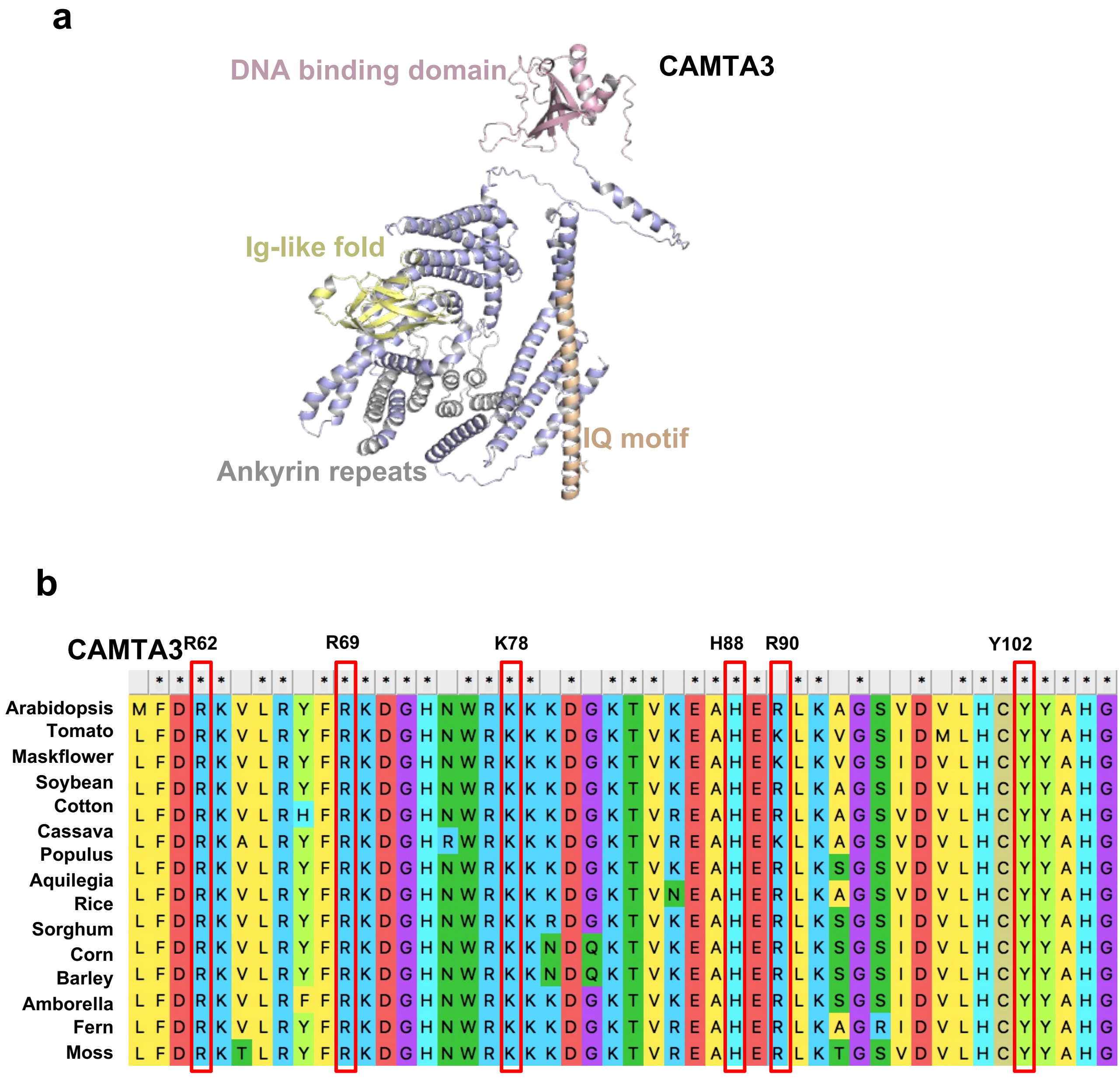


**Extended Data Figure 7. Predicted protein structure and conservation sites of CAMTA3.**
**a,** Predicted structural domain organization of CAMTA3 representing DNA binding domain (*light pink*), Immunoglobulin fold (*light yellow*), Ankyrin repeats (*light grey*) and Isoleucine/Glutamine (I/Q) motif. The structural analysis were based on highest model probability predicted by AlphaFold 3. **b,** Protein sequence alignment of CAMTA3 across representative land plant species shows the conservation of the R62, R69, K78, H88, R90 and Y102 site, the predicted amino acid interacting residues with RSER.

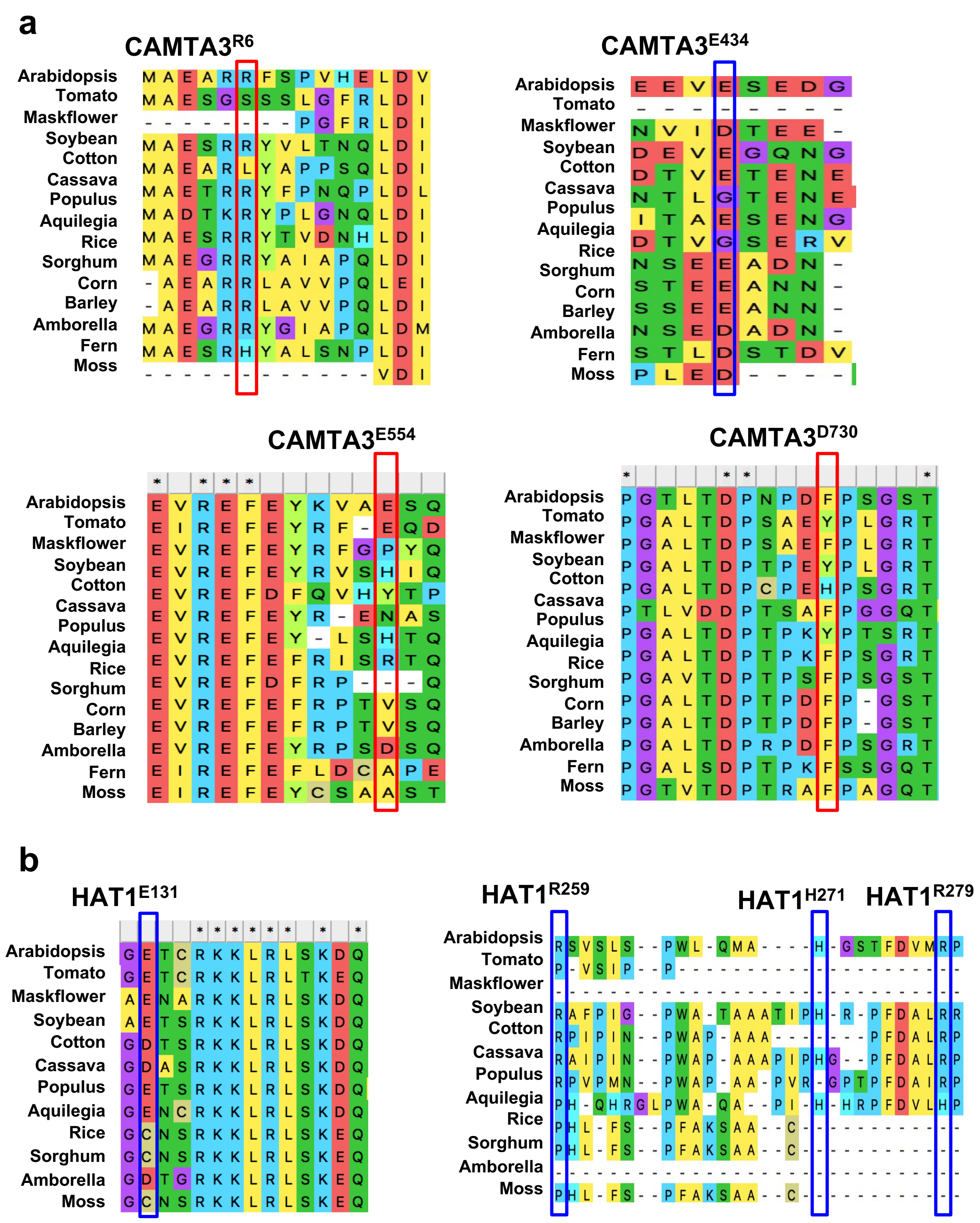


**Extended Data Figure 8. Evolutionary conservation of predicted CAMTA3-HAT1 interaction sites in land plants.**

**a,** Protein sequencing alignment of CAMTA3 across representative land plant species shows the conservation of the R6, E434, E554 and D730 site, which were predicted to be the interaction site with HAT1. **b,** Protein sequencing alignment of HAT1 across representative land plant species shows the conservation of the E131, R259, H271 and R279 site, which were predicted to be the interaction site with CAMTA3.

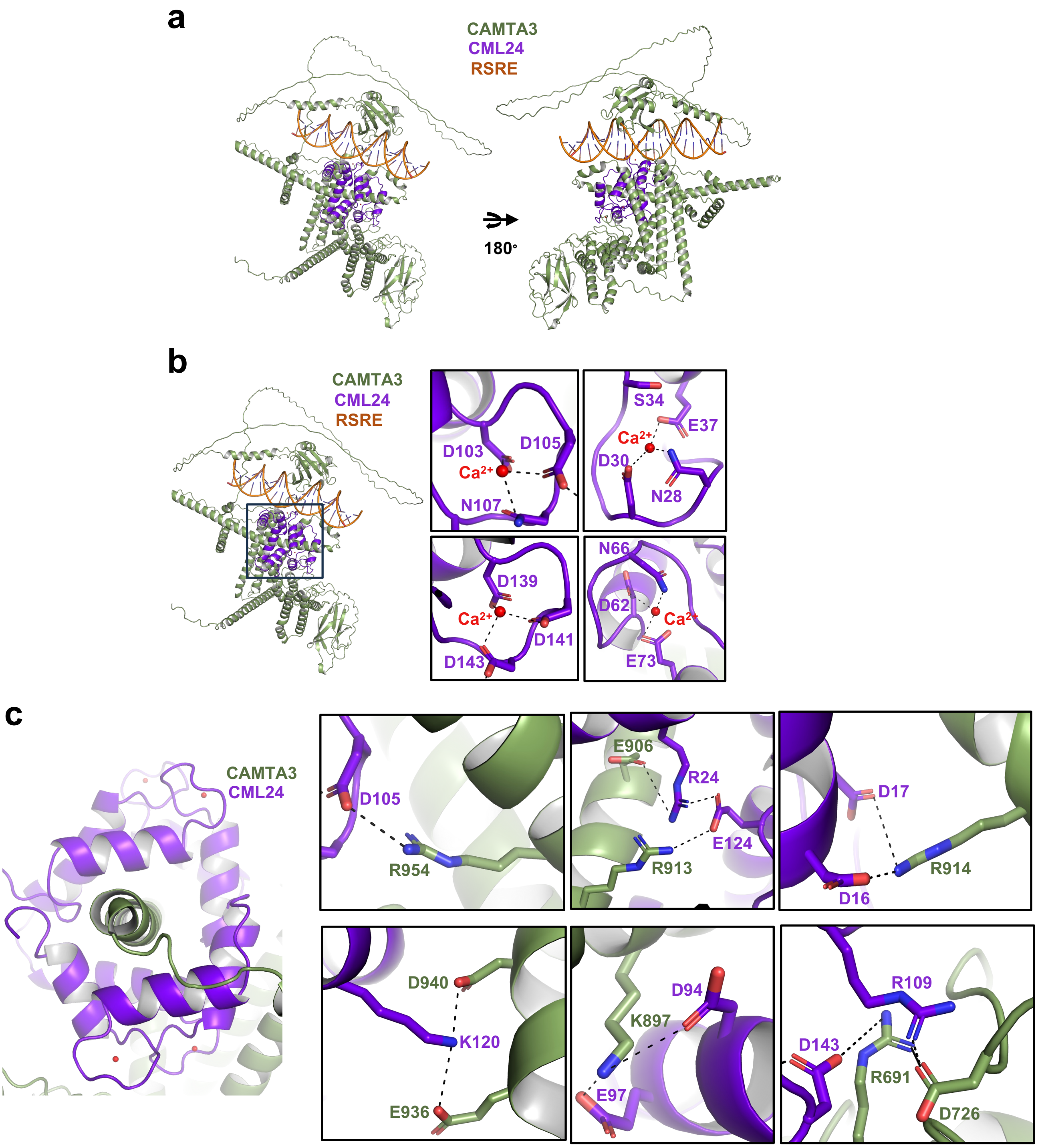


**Extended Data Figure 9. Predicted structure of RSRE-CML24-CAMTA3 complex.**
**a**, Structure of RSRE (orange)-CML24 (violet)-CAMTA3 (green) shown in different views. **b**, Stabilizing interaction of $Ca^{2+}$ (red sphere) bound to CML24 (left panel). Zoom-in view of Calcium (Ca2+, red sphere) stabilized by conserved aspartic acid residues. Dashed lines indicate co-ordination bond. **c**, Interaction of IQ motif of CAMTA3-CML24 (left panel). Zoom-in view of the interacting amino acids residues in the IQ motif of CAMTA3 with CML24 (right panel). Dashed lines indicate the polar salt bridges. All the structural analyses were based on the highest model probability predicted by AlphaFold 3.

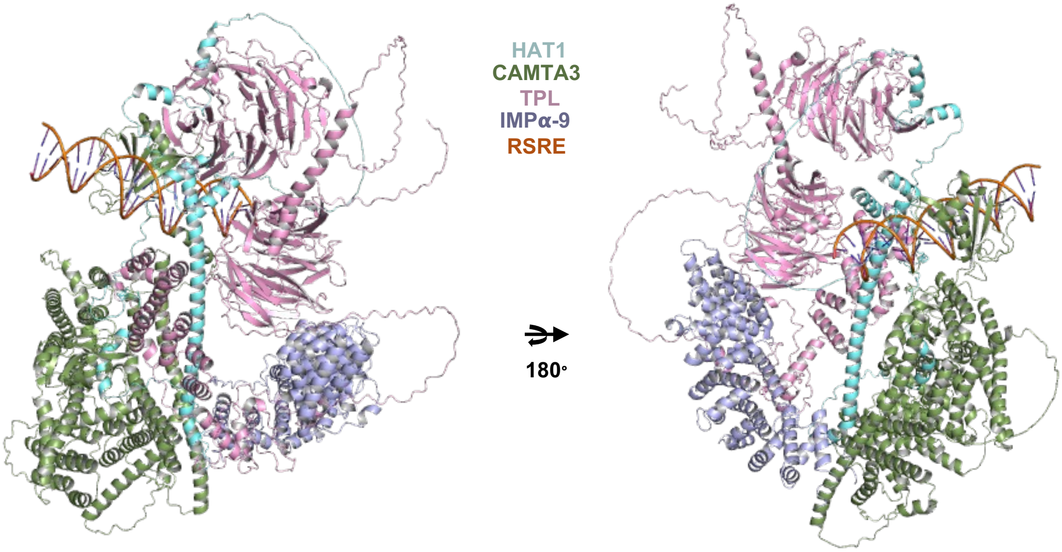


**Extended Data Figure 10. Predicted structure of RSRE-CAMTA3-HAT1-TPL-IMPα-9 complex.** Two views of structure complex of RSRE (orange)-HAT1 (blue)-CAMTA3 (green)-TPL (pink)-IMPα-9 (purple). All the structural analyses were based on the highest model probability predicted by AlphaFold 3.
